# Extracellular vesicles secreted by *Brugia malayi* microfilariae modulate the melanization pathway in the mosquito host

**DOI:** 10.1101/2022.04.11.487926

**Authors:** Hannah J. Loghry, Hyeogsun Kwon, Ryan C Smith, Noelle A Sondjaja, Sarah J Minkler, Sophie Young, Nicolas J Wheeler, Mostafa Zamanian, Lyric C Bartholomay, Michael J Kimber

**Affiliations:** Department of Biomedical Sciences, College of Veterinary Medicine, Iowa State University, Ames, Iowa, USA; Department of Entomology, College of Agriculture and Life Sciences, Iowa State University, Ames, Iowa, USA; Department of Pathobiological Sciences, College of Veterinary Medicine, University of Wisconsin-Madison, Madison, Wisconsin, USA

## Abstract

Vector-borne, filarial nematode diseases represent a significant and affecting disease burden in humans, domestic animals, and livestock worldwide. Parasitic filarial nematodes require both an intermediate (vector) host and a definitive (mammalian) host during the course of their life cycle. In either host, the nematode must evade the host elicited immune response in order to develop and establish infection. There is direct evidence of parasite-derived immunomodulation in mammals, however, there is less evidence of parasite immunomodulation of the vector host. We have previously reported that all life stages of *Brugia malayi*, a causative agent of lymphatic filariasis, secrete extracellular vesicles (EVs). Here we investigate the immunomodulatory effects of microfilariae derived EVs on the vector host *Aedes aegypti.* RNA-seq analysis of an *A. aegypti* cell line treated with *B. malayi* microfilariae EVs showed differential expression of both mRNAs and miRNAs, some with roles in immune regulation. One downregulated gene, AAEL002590, identified as a serine protease, was shown to have direct involvement in the phenoloxidase (PO) cascade through analysis of PO activity. Similarly, injection of adult female mosquitoes with *B. malayi* microfilariae EVs validated these results *in vivo*, eliciting a downregulation of the AAEL002590 transcript and a significant reduction in PO activity. Our data indicates that parasite-derived EVs are capable of interfering with critical immune responses in the vector host, particularly immune responses such as melanization that target extracellular parasites. In addition, this data provides novel targets for transmission control strategies for LF and other parasitic diseases.

**Author Summary:** Vector-borne, filarial nematode diseases represent a significant and affecting disease burden in humans, domestic animals and livestock worldwide. Parasitic nematodes must evade the elicited immune response of their hosts in order to develop and establish infection. While there is evidence for immunomodulation of the mammalian host, the mechanism of this immunomodulation is not fully clear and there is limited evidence for immunomodulation of the vector host. Here we have shown that parasite-derived extracellular vesicles are effector structures for immunomodulation of the vector host. In particular, we have identified that parasite-derived extracellular vesicles can interfere with critical mosquito immune responses against parasites. This data provides insight into parasite biology and novel targets for transmission control strategies for parasitic diseases.

## 1. Introduction

Vector-borne, filarial nematode diseases represent a significant and affecting disease burden in humans, domestic animals, and livestock worldwide. In humans, Lymphatic Filariasis (LF) is caused by multiple species of filarial nematodes, including *Brugia malayi* and is endemic in 72 countries with over 860 million people infected or at risk of infection (1). Adult parasites reside in the lymphatic vasculature and although often asymptomatic, infection can result in extreme morbidity including lymphangitis, lymphedema (primarily in the extremities), and secondary bacterial infection/dermatitis (2). Current control strategies rely on mass drug administration programs that utilize inadequate anthelmintic drugs that do not effectively kill adult parasites or resolve established infections. The need for new control strategies of filarial nematode diseases is necessary, however, progress in developing effective treatments has been stalled by our lack of understanding of parasite biology and host-parasite interactions.

Parasitic filarial nematodes require both an intermediate (vector) host and a definitive (mammalian) host during the course of their life cycle. In either host, the nematode must evade the elicited immune response of the host in order to develop and establish infection. Various immune evasion strategies have been documented, including manipulation of host immune responses (3). In mammals, there is direct evidence of parasite-derived immunomodulation. It has been shown that parasites are capable of expanding regulatory immune cells (4–10), inducing apoptosis in type 1 immune response cell types (11–14), manipulating pattern recognition receptors (PRRs) (15–21), and increasing anti-inflammatory cytokines such as IL-4/IL-10 (22–24). However, there is less extensive evidence of parasite immunomodulation of the vector host. Early studies have shown that filarial nematode parasites can inhibit melanization (25).

Melanotic encapsulation, a crucial mosquito innate immune response to the microscopic larval stages of the parasite that infect mosquitoes, is a core arthropod defense mechanism that prevents infecting nematode growth and reproduction, and eventually leads to their death (26). Melanotic encapsulation involves both the humoral and cellular components of the innate immune response in mosquitoes. Upon recognition of a pathogen, the cellular arm of insect innate immunity drives aggregation of hemocytes to form a multicellular layer around the invading pathogen.

Concurrently, the humoral arm of insect innate immunity initiates melanin production in the hemocytes (27–35). This process is controlled by the phenoloxidase (PO) cascade, which is initiated when a pathogen associated molecular pattern (PAMP) binds to its pattern recognition receptor (PRR) to initiate a serine protease cascade. This cascade ultimately leads to the activation of a pro-phenoloxidase activating factor which in turn will activate phenoloxidase (26,36,37). Phenoloxidase can then oxidize phenols to quinones which are further polymerized to melanin (38). Death of the parasite is believed to be due to nutrient deprivation, asphyxiation, and/or through the production of toxins such as quinones and other reactive oxygen species produced during melanin production (39, 40).

The mechanistic basis for nematode manipulation of mosquito immune responses is not clear but recent studies exploring vector host global transcriptomic changes in response to parasite invasion have identified downregulation of immune-related genes during infection (41–43). A consensus view is that the cumulative effect of this modulation, be it within the intermediate or definitive host, is to suppress the host immune response towards a tolerant state in which the immune response is still present and active, but damage to the parasite is limited. While there is unequivocal evidence that parasites can directly modulate the host immune response, and although the concept is broadly accepted, the parasite-derived effectors that drive this modulation at the cellular and molecular level remain unclear and poorly understood, especially within the context of the vector host.

Parasite excretory-secretory products (ESP) are a well-established source of potential effector molecules. Parasite ESP encompass freely secreted proteins and nucleic acids, as well as extracellular vesicles (EVs), which are membrane-bound structures secreted by both prokaryotic and eukaryotic cells including filarial nematodes (15,44–47). They contain complex cargo that can include proteins, small RNA species, and lipids (48, 49) and have been shown to be highly involved in cell-to-cell communication and have roles in various physiological processes (48,50– 52). Although EVs are a newly recognized fraction of parasitic nematode ESP, the cargo of some nematode EVs have been profiled, revealing contents to include protein and small RNA species with predicted immunomodulatory properties (15,44,46,53–61). There is strong evidence that these EVs have direct involvement in immunomodulation of mammalian hosts (4,15,16,44,53,55,56,60,62,63).

We hypothesize that filarial nematode EVs secreted by infective stages of filarial parasites act as effectors to modulate the immune response of the vector host mosquito. To test this hypothesis, we examined the modulatory effects of *B. malayi* microfilariae (mf) EVs on the global transcriptomic profile of *Aedes aegypti* derived Aag2 cells, an established model for mosquito hemocytes due to their characterized immunocompetence (64). We found that nematode EV treatment drove differential expression of host genes, including a serine protease gene. This gene was shown to have direct involvement in the PO pathway as knockdown of the gene lead to a reduction in PO activity *in vitro.* The effect of microfilariae EVs was subsequently investigated *in vivo,* and it was found that these microfilariae derived EVs inhibited PO activity in adult female *A. aegypti*. These findings provide evidence that parasite derived extracellular vesicles contain cargo that are capable of modulating critical vector host immune responses.

## 2. Results

### 2.1 *B. malayi* mf derived EVs are internalized by Aag2 cells

To confirm that EVs were being isolated from spent media, EVs were imaged using TEM (Fig. 1A). Particles isolated from spent media exhibited the classic exosome-like deflated soccer ball morphology under EM but such structures were absent from unconditioned media. Vesicle size was further validated with nanoparticle tracking analysis using NanoSight LM10 (Malvern Panalytical, Malvern UK)(Fig. 1B) and showed that the isolated EVs had a mean size and concentration of 92.2 nm and 2.68 x 10^9^ particles/ml respectively, well within the expected 50-200nm range. To investigate the potential for parasitic excretory-secretory products to interact with vector host immune cells, we treated Aag2 cells, an immunocompetent *A. aegypti* cell line (64), with PKH67 stained *B. malayi* mf derived EVs. 24 hours after treatment, cells were additionally stained with DAPI and phalloidin, and imaged with confocal microscopy. Aag2 cells were shown to internalize *B. malayi* mf derived EVs as compared to control cells (Fig. 1C-D).

**Figure 1.**
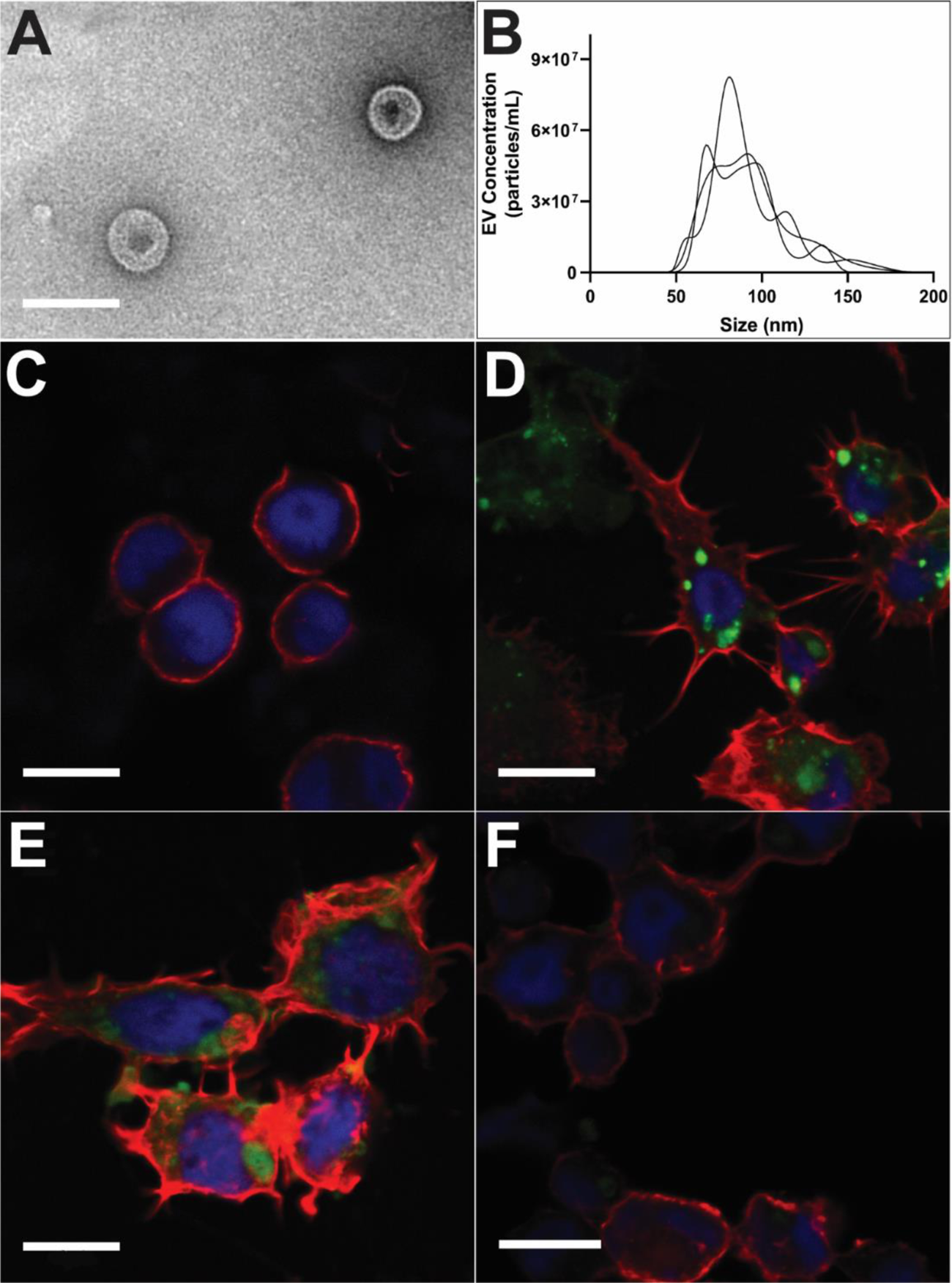
*B. malayi* mf derived EVs are internalized by Aag2 cells. Isolation of *B. malayi* mf EVs was confirmed by TEM (A) and size profile was further validated with nanoparticle tracking analysis (B). PKH67 stained *B. malayi* mf EVs were incubated with Aag2 cells for 24 hours. Cells were stained with Alexa Fluor 647 Phalloidin and DAPI and imaged with a Leica SP5 X MP confocal/multiphoton microscope system. 51% of cells incubated with PKH67 stained EVs showed internalization indicated by the green fluorescence inside the cell (D) as compared to control cells (C). Cells treated with endocytosis inhibitors chlorpromazine (E) showed no endocytosis of stained EVs while cells treated with nystatin (F) showed diffuse uptake of EVs throughout the cytoplasm. Scale bar (A) = 150 nm. Scale bar (C-F) = 10 µM.

EVs appeared in punctate areas within the cell and were not found diffused throughout the cytoplasm. This correlates with previous evidence that EVs are internalized via endocytosis and thus would be confined to endosomes within the cytoplasm (65). In addition, internalization of parasite EVs by murine epithelial cells showed a similar punctate appearance (15). However, a different phenotype was seen by parasite EVs internalized by murine macrophages and human monocytes where the EVs appeared diffused throughout the cytoplasm (44,46,53). These differences in internalization appearance may be due to various endocytosis pathways utilized by the various cell types. To begin to tease apart the endocytic mechanism by which mf EVs are being internalized, Aag2 cells were treated with the endocytosis inhibitors chlorpromazine (CPZ) and nystatin. CPZ is an inhibitor of clathrin-mediated endocytosis and has been shown to inhibit the function of a key clathrin-mediated endocytic adaptor protein AP2 (66, 67). Nystatin is capable of binding cholesterol and thus can inhibit caveolin-mediated endocytosis (68). It was observed that chlorpromazine, but not nystatin inhibited the endocytosis of *B. malayi* mf EVs (Fig. 1E-F), suggesting that the mechanism of endocytosis of parasitic EVs is clathrin-mediated. EV internalization was quantified using flow cytometry (Supplemental Fig. 1). 51% of Aag2 cells internalized *B. malayi* mf EVs as compared to untreated cells (p < 0.0001, N = 3). Treatment with the endocytosis inhibitor chlorpromazine reduced the number of Aag2 cells that internalized *B. malyi* mf EVs by 39% as compared to EV only treated cells (p = 0.0003, N = 3). However, the endocytosis inhibitor nystatin did not significantly inhibit EV internalization as compared to EV only treated Aag2 cells.

### 2.2 EV Treatment suppresses miRNA expression with Immune Related Targets

Due to the immunomodulatory cargo identified in other *B. malayi* life stages (44, 46), we hypothesized that Aag2 cell phenotypes would be modulated by treatment with *B. malayi* mf EVs. To simulate a naturally occurring infection, Aag2 cells were first treated with LPS (500ng/ml) to mimic the immune response that would initially occur during the early stages of infection. 12 hours later, the cells were then treated with either dPBS or *B. malayi* mf EVs to examine the modulatory effects of EV treatment on an established response. 16 hours later cells were collected and processed for miRNA sequencing. Of the 300 miRNAs identified, 196 were expressed in all three treatment groups (control, LPS only, and LPS + EV). The control treatment group shared 21 miRNAs with the LPS only treatment group and 12 with LPS + EV while LPS and LPS + EV shared 10 common miRNAs. The control, LPS and LPS + EV treatment groups had 40, 19 and two miRNAs that were unique to each treatment group, respectively (Fig 2A). To investigate the ability of *B. malayi* EVs to regulate an immune response, we compared miRNA expression between LPS and LPS + EV treatment groups. Six miRNAs were identified to be significantly downregulated in LPS+EV as compared to LPS only, including aae-mir-1175, aae-mir-2945, bmo-mir-6497, nlo-mir-275, aae-mir-184, and PC-5p-30141_33 (Fig 2B). Target prediction was conducted on these differentially expressed miRNA followed by GO analysis of the predicted gene targets. Targets were identified for five out of the six downregulated miRNAs with gene targets of these downregulated miRNAs having roles in proteolysis, regulation of transcription, signal transduction, phagocytosis, and cell differentiation among others (Fig 2C).

**Figure 2.**
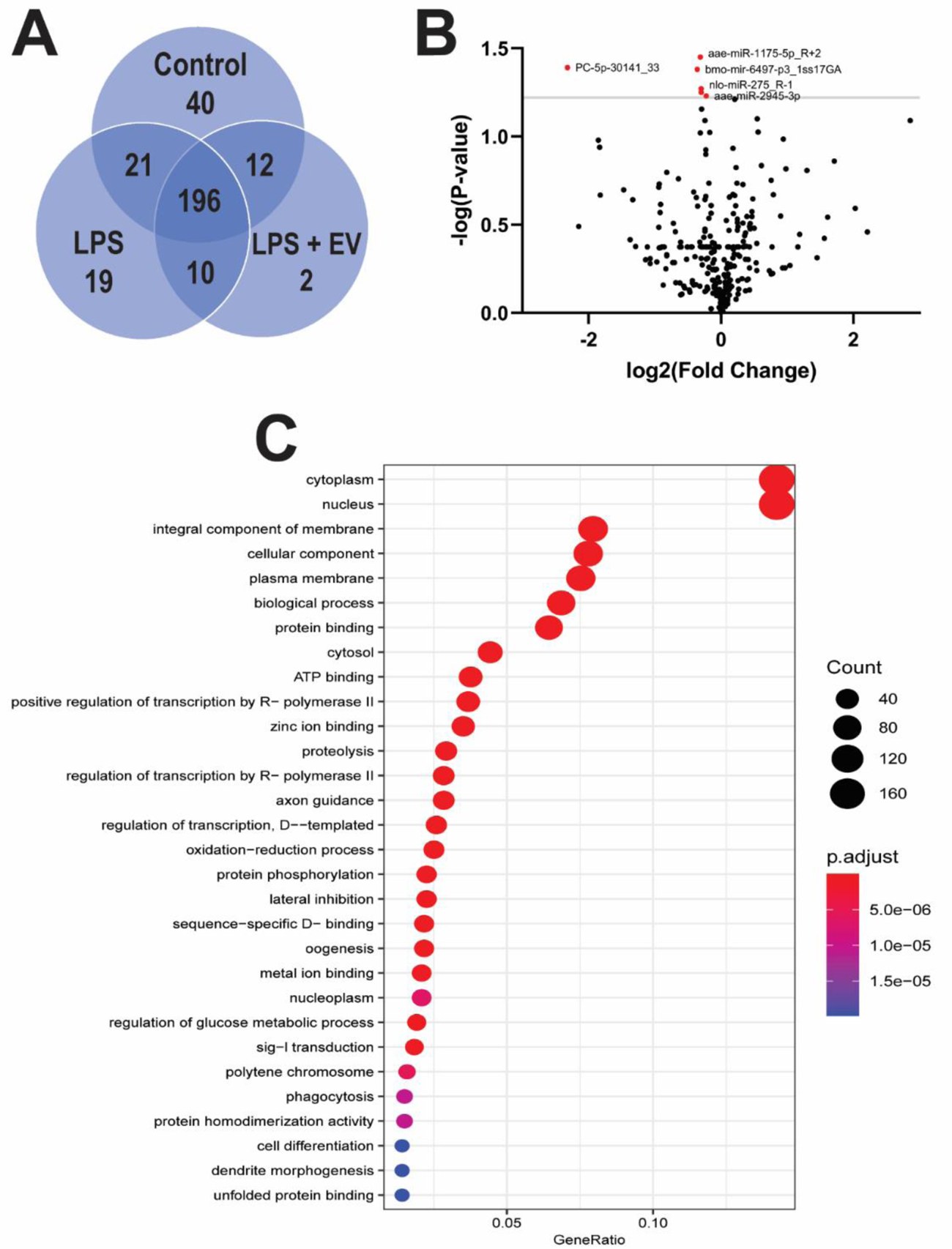
EV Treatment suppresses miRNA expression with Immune Related Targets. miRNA-seq analysis was performed on control, LPS and LPS + EV treated Aag2 cells. All three treatment groups shared 196 miRNAs while 40, 19 and two miRNAs were unique to control, LPS only and LPS + EV treatment groups respectively (A). Six significantly, differentially expressed miRNAs were identified between the LPS and LPS + EV treatment groups (B). Predicted targets were identified for five out of the six significantly downregulated miRNAs. Gene ontology (GO) analysis of these predicted gene targets identified their role in various physiological processes including proteolysis, signal transduction and regulation of transcription (C).

Additionally, KEGG analysis identified that the predicted gene targets are involved in multiple immune related pathways (Table 1). Gene targets of the downregulated miRNAs are predicted to be involved in common insect immune signaling pathways such as Toll/IMD, MAPK, TGFβ and insulin signaling pathways among others. Some of the predicted gene targets include AAEL008634, a jnk protein; AAEL010433, a transcriptional co-repressor, AAEL003505; a jun protein; and AAEL013433, a spaetzle-like cytokine. One of these miRNAs, aae-mir-1175, is conserved in *Anopheles gambiae* and was shown to be downregulated in plasmodium infected mosquitoes as compared to non-infected mosquitoes (69). In addition, mir-1175 has been identified to be solely expressed in the mosquito midgut a critical barrier in parasite development and transmission in the vector host. A similar phenotype was seen in *A. aegypti* where aae-mir-1175 was downregulated in mosquitoes infected with dengue virus as compared to non-infected mosquitoes(70). This provides evidence for a conserved immunomodulation phenotype across diverse vector pathogens that enables pathogen migration and development. *A. aegypti* infected with *Wolbachia* showed a similar downregulation of aae-mir-2945 when compared to non-infected mosquitoes providing additional support for downregulation of miRNAs to drive immunomodulation. The direct role that downregulating these miRNAs have on insect immune cell responses remain unknown, but these data suggest that *B. malayi* EVs are capable of modulating post-transcriptional control of host gene expression, including genes potentially involved in mosquito immune signaling pathways. Additional experimentation needs to be conducted to determine what specific effector molecule in the EV cargo is driving this modulation.

**Table 1.** KEGG Analysis of downregulated miRNA predicted targets KEGG analysis of the predicted target genes of the significantly, downregulated miRNAs revealed an enrichment of immune signaling pathways, including common insect immune signaling pathways such as Toll/IMD, MAPK, TGFβ and insulin signaling.

| miRNA | Function of Predicted Target |
| --- | --- |
| aae-mir-1175 | MAPK Signaling Pathway |
|  | Longevity Regulating Pathway |
|  | Phosatidylinositol Signaling Pathway |
|  | NOD-like Receptor Signaling Pathway |
|  | Wnt Signaling Pathway |
|  | Notch Signaling Pathway |
| | TGFβ Signaling Pathway |
|  | FoxO Signaling Pathway |
|  | Toll and Imd Signaling Pathway |
| aae-mir-2945 | Phosatidylinositol Signaling Pathway |
|  | MAPK Signaling Pathway |
|  | Wnt Signaling Pathway |
|  | Toll and Imd Signaling Pathway |
|  | Insulin Signaling Pathway |
|  | Neurotrophin Signaling Pathway |
| | TGFβ Signaling Pathway |
|  | FoxO Signaling Pathway |
|  | Phosatidylinositol Signaling Pathway |
|  | mTOR Signaling Pathway |
|  | Longevity Regulating Pathway |
|  | Ras Signaling Pathway |
| bmo-mir-6497 | Toll and Imd Signaling Pathway |
|  | MAPK Signaling Pathway |
|  | Notch Signaling Pathway |
|  | Longevity Regulating Pathway |
|  | mTOR Signaling Pathway |
|  | Wnt Signaling Pathway |
|  | FoxO Signaling Pathway |
|  | p53 Signaling Pathway |
|  | Hedgehog Signaling Pathway |
|  | Rap1 Signaling Pathway |
|  | Insulin Signaling Pathway |
|  | Neurotrophin Signaling Pathway |
| PC-5p-30141_33 | Hedgehog Signaling Pathway |
|  | MAPK Signaling Pathway |
| | TGFβ Signaling Pathway |
|  | mTOR Signaling Pathway |

### 2.3 Microfilariae EVs downregulate predicted immune related genes *in vitro*

mRNA-seq analysis was conducted concurrently with miRNA-seq analysis. Many differentially expressed genes between LPS and LPS + EV treatment groups were identified (Fig. 3A). A majority of the most highly upregulated or downregulated genes were uncharacterized protein coding genes with unknown function. Thus, rather than focusing on these most differentially regulated targets, genes that were significantly differentially expressed but also moderately annotated were instead chosen for *in vitro* validation. For example, AAEL024490 is a predicted cys-loop ligand-gated ion channel (cysLGIC) subunit with high sequence identity to a predicted gamma-aminobutyric acid (GABA) gated chloride ion channel (CLIC) subunit (this subunit will be referred to as a CLIC subunit for simplicity). AAEL002590 is a putative serine protease that has a *Culex quinquefasciatus* ortholog that has been identified as a pro-phenoloxidase activating factor (PPAF). Both the CLIC subunit gene and the serine protease gene were significantly downregulated upon EV treatment by 99% (p = < 0.0001) and were among the genes chosen for *in vitro* validation using RT-qPCR. Aag2 cells were stimulated with LPS to elicit an immune response and then followed with treatment of serial dilutions of *B. malayi* mf EVs. The CLIC subunit was significantly downregulated, expression was reduced by 68% when treated with 1×10^5^ EVs (p = 0.0369, N = 3) (Fig. 3B) as compared to LPS only treated cells. EV treatment suppressed CLIC expression to basal levels observed in non-LPS treated Aag2 cells (53% of LPS stimulated value, p = 0.0425, N = 3). The serine protease gene was also significantly downregulated after treatment with 1×10^5^ *B. malayi* mf EVs (57%, p = 0.0223, N = 3) (Fig. 3C). Again, EV treatment completely abrogated the LPS stimulation of expression (p = 0.0271, N = 3). These findings are biologically relevant as 1×10^5^ EVs is within the range of anticipated EVs that would be present in a mosquito after a blood meal. While the number of mf taken up by a mosquito during a blood meal varies on the microfilariae density in the blood of the host, it has been established that the approximate mean number of mf taken up by a mosquito is between 1-300 mf with most taking up approximately 40 mf (71–73). In addition, it has been shown that *B. malayi* mf secrete, on average, 4000 EVs per mf in 24 hours (47). These data provide the approximate range of EVs that would be present in a mosquito within 24 hours of a blood meal would be between 1 x 10^5^ – 1 x 10^6^. Since AAEL002590 was identified as a serine protease with homology to a *C. quinquefasciatus* PPAF, we next wanted to investigate whether this gene was involved in the PO pathway. RNAi was used to knockdown AAEL002590 in Aag2 cells with a time course experiment showing that optimal knockdown occurred at 24 hrs post-RNAi treatment with 79% suppression of AAEL002590 expression (p = 0.0012, N =3) (Supplemental Figure 3). To investigate whether AAEL002590 was involved in the PO pathway, Aag2 cells were treated with duplexed siRNA or scrambled siRNA as a negative control for 24 hrs.

**Figure 3.**
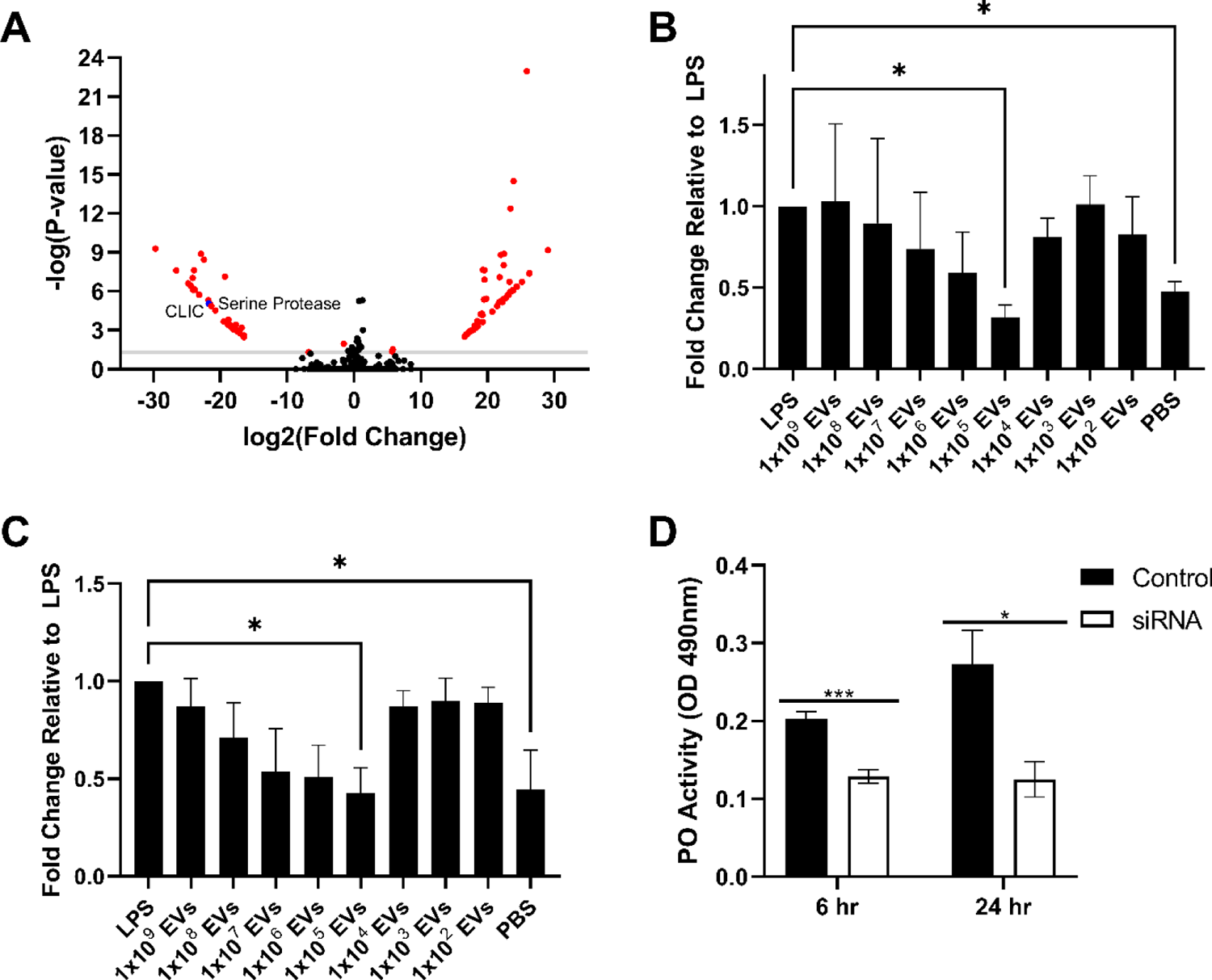
EVs released by *B. malayi* microfilariae downregulate predicted immune related genes *in vitro*. Multiple genes were differentially expressed between LPS and LPS + EV treatment groups (A). Two moderately annotated and significantly downregulated genes were chosen for further *in vitro* validation by RT-qPCR. Both the CLIC subunit gene (B) and the serine protease gene (C) were significantly downregulated when treated with 1×10^5^ *B. malayi* mf EVs as compared to control. RNAi knockdown of the serine protease gene in Aag2 cells inhibited phenoloxidase activity as compared to control at both 6 and 24 hrs post treatment (D) indicating that the serine protease gene is involved in the PO pathway. N = 3 (minimum). Mean ± SEM. * P < 0.05, ***P < 0.001.

Following RNAi treatment cells were either treated with dPBS to quantify changes in basal PO activity or challenged with LPS (500ng/ml) for 6 or 24 hours after which cell culture supernatant was collected and mixed with L-DOPA for the PO activity assay. The assay was incubated overnight and basal PO activity was measured at 490nm. Basal PO activity was inhibited by 36% after AAEL002590 RNAi as compared to control cells at 6 hours (p = 0.0002, N = 3) and inhibited by 54% as compared to control at 24 hours (p = 0.018, N = 3) (Fig 3D). While LPS treatment has been used to induce an immune response in Aag2 cells previously (64) and was successful in inducing an immune response in Aag2 cells as evident by our gene expression experiments, LPS did not sufficiently induce the PO cascade *in vitro*. However, it is clear from RNAi-mediated knockdown of basal PO activity that AAEL002590, a target for parasite EV modulation, is involved in the host PO pathway.

GO enrichment analysis was conducted on all significantly (p ≤ 0.05) upregulated or downregulated mRNAs. We found that genes upregulated following EV treatment were enriched for GO terms associated with metabolic processes and oxidoreductase activity (Fig. 4A). Some increases in metabolic activity and increases in oxidoreductase activity can be explained by the vector’s reaction to initial parasite infection. However, increases in steroid and lipid biosynthesis may be driven by parasite effector molecules. It has been shown that host steroid hormones can be influenced by development of parasites and dictate their course of infection, with increase production of steroid hormones leading to more rapid development and longer infections (74, 75). In addition, it has been shown that host lipid biosynthesis is hijacked by parasites and is a common them in vector-borne diseases(76).

**Figure 4.**
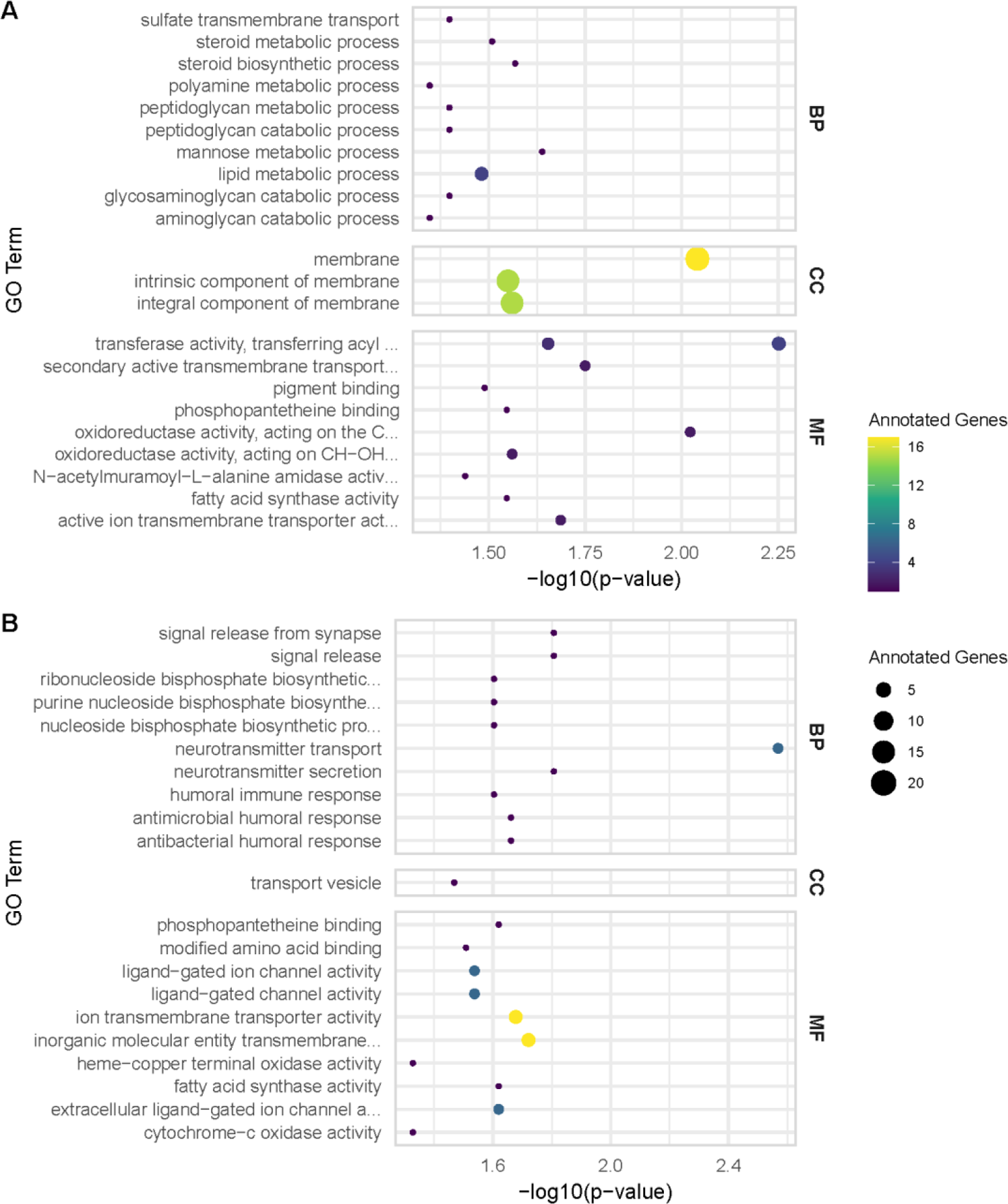
Downregulated mRNAs are involved in signaling and immune responses. GO analysis on significantly upregulated genes (A) shows that these genes are enriched in GO terms associated with metabolic processes and oxidoreductase activity while downregulated genes (B) are enriched for GO terms associated with signaling and immune responses.

Genes that were downregulated after EV treatment were enriched for GO terms associated with signaling and immune responses (Fig. 4B). Signaling GO terms include ligand-gated ion channel activity, transmembrane ion transporter activity, neurotransmitter release and neurotransmitter secretion. As mentioned previously, we have already validated that a predicted GABA-gated chloride ion channel subunit is downregulated after EV treatment. The GABAergic system has been highly implicated in human immune functions including roles in phagocytosis, cytokine production, and cell proliferation(77). While the main findings in humans have concluded that the GABAergic system leads to immunosuppressive phenotypes, the possible role of GABA receptors in invertebrate immune responses has not been well studied. In insects, the cysLGIC superfamily is known for its inhibitory roles in neurotransmission and as target sites for insecticides(78). Dieldrin and endosulfan are organochlorine-based insecticides that function as GABA receptor antagonists and it has been shown that sub-lethal doses of both dieldrin and endosulfan inhibited the encapsulation of *Leptopolina boulardi* eggs by *Drosophila* larvae(79). The strong body of evidence that GABA receptors are involved in mammalian neurohormonal immune regulation and that certain GABA receptor antagonists in insects can modulate encapsulation, suggests a potential role for the predicted GABA-gated chloride channel in neurohormonal regulation of mosquito immune responses. In particular, a role in promoting or driving the encapsulation process.

### 2.4 Phenoloxidase activity is inhibited by EV treatment

Having established that parasite EV treatment modulates AAEL002590 in Aag2 cells *in vitro*, we next wanted to determine if this phenotype was recapitulated *in vivo*. Adult female mosquitoes were injected with LPS (1mg/ml) followed by injection with mf EVs or dPBS 6 hours later. Mosquitoes were incubated for 24 hours and then AAEL002590 expression was assayed by RT-qPCR. Injection with 1×10^5^ mf EVs significantly downregulated the serine protease gene by 84% (p = 0.02, N = 3) as compared to LPS only (Fig. 5A). This EV-suppression returned AAEL002590 expression to basal levels comparable to control mosquitoes in which AAEL002590 expression was 74% lower than LPS stimulated mosquitoes (p = 0.05, N =4). Since we had already shown that knockdown of the serine protease gene in Aag2 cells inhibited PO activity *in vitro* we wanted to investigate if *B. malayi* mf EVs could inhibit PO activity *in vivo.* Adult female mosquitoes were treated as previously described and hemolymph was collected by perfusion following the 24-hour incubation. Hemolymph was then mixed with L-DOPA and PO activity was measured by optical density (OD) readings at 490nm every 5 minutes for 30 minutes and a final reading at 60 minutes. PO activity was significantly inhibited in hemolymph from mosquitoes injected with 1×10^5^ mf EVs at all time points. Specifically, PO activity was inhibited by 65% (p < 0.05), 81% (p < 0.0001), 80% (p < 0.0001), 78% (p < 0.0001), 76% (p < 0.0001), 74% (p < 0.0001), 72% (p < 0.0001), and 64% (p < 0.0001) at 0, 5, 10, 15, 20, 25, 30 and 60 minutes respectively (all N = 3) as compared to LPS only treated mosquitoes. These results indicate that *B. malayi* mf EVs inhibit PO activity *in vivo* at biologically relevant concentrations. While the PO activity induced by LPS treatment may not have appeared as high as predicted, the pronounced inhibition after EV treatment is compelling.

**Figure 5.**
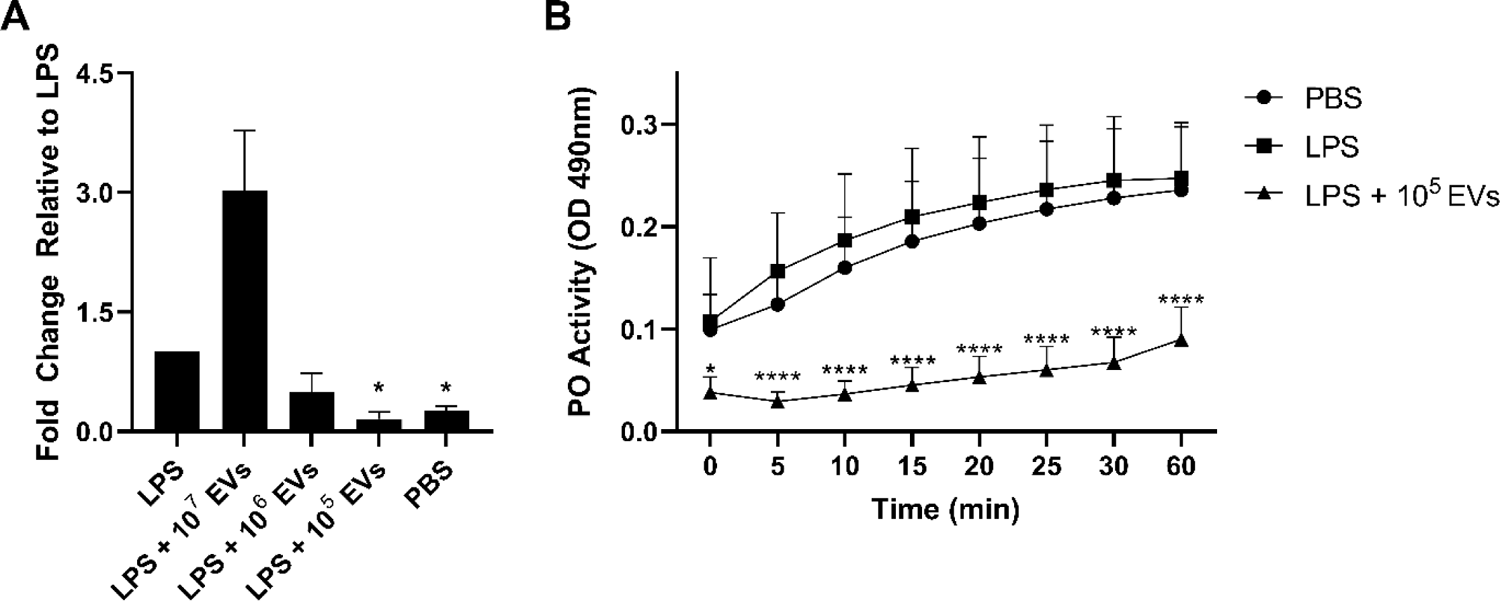
Phenoloxidase activity is inhibited by EV treatment. Validation of the downregulation of the serine protease gene *in vivo* was investigated by injection of adult female mosquitoes with serial dilutions of *B. malayi* mf EVs after initial treatment with LPS. 1×10^5^ EVs significantly downregulated the serine protease gene as compared to LPS only (A). Hemolymph of injected mosquitoes was collected to test for phenoloxidase activity. Treatment of adult female mosquitoes with 1×10^5^ mf EVs inhibited PO activity as compared to LPS only at all time points (B). N = 3 (minimum). Mean ± SEM. *P < 0.05, ****P < 0.0001.

## 3. Discussion

While there is strong evidence for parasite-derived host immunomodulation of the mammalian host, evidence of immunomodulation of the vector host is lacking. Here we have shown that parasite-derived extracellular vesicles (EVs) elicit transcriptional changes in an insect immune cell model, specifically, our data show that *B. malayi* mf derived EVs can modulate multiple genes involved in the humoral immune response. The humoral immune response of a mosquito is comprised of pattern recognition receptors (PRRs), antimicrobial peptides (AMPs) and components of the phenoloxidase (PO) cascade. Downregulation of genes involved in these immune responses would be advantageous for any invading pathogen, especially those that must migrate through the mosquito hemolymph. Melanotic encapsulation is a fundamental mosquito defense mechanism against parasites that involves the PO cascade and here we have identified a serine protease with homology to a known prophenoloxidsae activating factor (PPAF) that is downregulated when mosquito cells are treated with *Brugia* EVs. Independent RNAi-mediated knockdown of this serine protease leads to an inhibition in PO activity in an insect cell line. We were also able to show that this phenotype is recapitulated during *Brugia* infection of mosquitoes *in vivo.* Further experimentation is needed to determine if this serine protease is indeed a true PPAF or if it is a serine protease involved in an upstream cascade that activates pro-PPAF. In either case, however, our data provides evidence that parasite-derived EVs are effector structures in immunomodulation of vector hosts with the ability to interfere with critical host immune responses (Fig 6). Modulation of the vector host melanization immune response is logical as it is the main vector defense mechanism against large, extracellular pathogens such as parasitic nematodes. While our data provides novel mechanistic evidence for modulation of the host melanization immune response, this phenomenon seems to be central to parasite-vector host interactions. Christensen and LaFond (1986) were the first to provide evidence for parasite-derived modulation of the melanization response, showing that *B. pahangi* infected *A. aegypti* had reduced ability to melanize when challenged with intrathoracic inoculation of new *B. pahangi* mf (25). In addition, targeting of the melanization and encapsulation immune response is a common phenotype seen in infections of *Galleria mellonella* with the parasitic nematode *Steinernema carpocapsae.* Studies have shown that a trypsin-like serine protease secreted by *S. carpocapsae* can inhibit PO activity *in vitro* and affects the morphology of *S. capocapsae* hemocytes and inhibits their ability to spread, a feature necessary for encapsulation (80). Further, a secreted chymotrypsin protease from *S. carpocapsae* has also been shown to inhibit PO activity and encapsulation of *G. mellonella* hemocytes both *in vitro* and *in vivo* (81). *Brugia* are known to actively secrete a number of proteases some of which may be involved in modulating the melanization response; indeed, a cathepsin L-like protease is abundantly found in the EVs of infective third stage larvae isolated from *A. aegypti* (44) that is essential to parasite survival within the mosquito (82). An important next step will be to characterize the cargo of *B. malayi* mf EVs to identify those effector molecules responsible for PO pathway downregulation. As this work continues, it will be essential to consider that the modulatory molecules may not be proteins. We have shown that filarial nematode EVs also contain a diverse miRNA cargo (44) and secreted EVs represent a way that effector miRNAs can be released from the parasite and protected during trafficking to host cells, where they might downregulate immune pathways at the genetic level.

**Figure 6.**
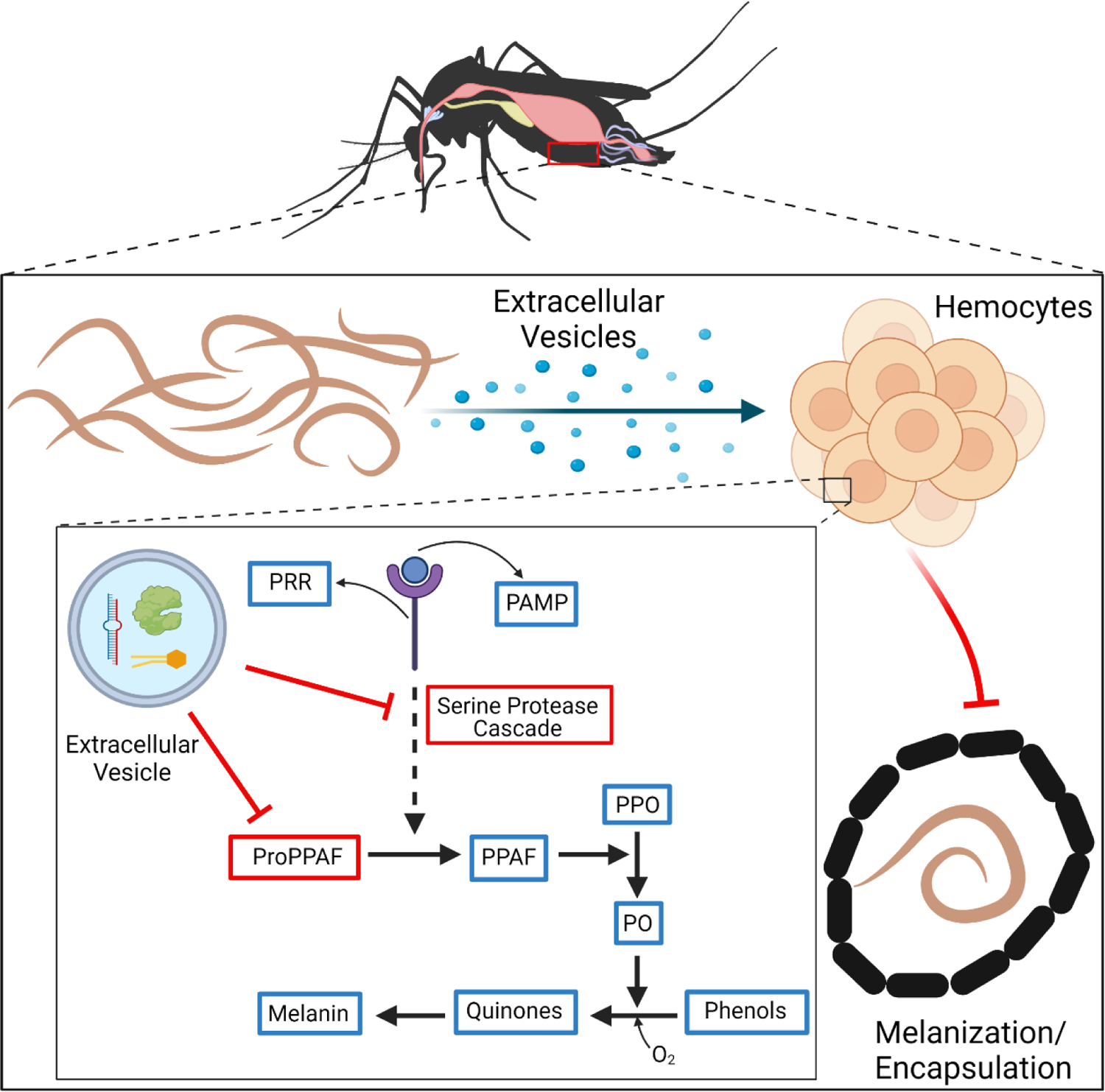
*B. malayi* microfilariae release EVs that interfere with the PO cascade and melanization. Melanotic encapsulation is a common insect defense mechanism against parasites. Upon recognition of a parasite, hemocytes aggregate forming a multicellular layer that deposits a melanin-enriched capsule around the invading parasite. Melanin production is controlled by the phenoloxidase (PO) cascade, which through a series of interdependent reactions, leads to the activation of PO that oxidizes phenols to quinones, which are further polymerized to melanin. Death of the parasite is believed to be due to nutrient deprivation, asphyxiation, or through the production of toxins such as quinones and other reactive oxygen species produced during melanin production. *B. malayi* microfilariae-derived extracellular vesicles downregulate a serine protease that functions either at the serine protease cascade or as a PPAF, either way interfering with the production of PO and thus inhibiting melanization of invading parasites.

To this end, several transcriptomic studies have looked at the global transcriptional changes that occur in the mosquito host during parasite infection (41–43,83–85). Many of these studies also identified parasite-derived downregulation of mosquito host serine proteases at the genetic level. A study conducted on *B. malayi*-infected *Armigeres subalbatus* showed that there was a significant reduction in expression of multiple serine protease genes during the first 24 hours of infection (41). This correlates with the time frame that EV-secreting mf would be migrating from the midgut, through the hemocele and to the thoracic musculature, and the timeframe seen in our studies. While *B. malayi* do not effectively develop to infective L3 stage parasites in *A. subalbatus*, it still provides evidence for host transcriptional changes during early stages of infection. Similar trends were observed in *B. malayi* infected *A. aegypti* where there was evidence for parasite derived alteration in expression of genes involved in blood digestion and immune function including specific downregulation of serine protease genes (43), providing broad evidence for parasite immunomodulation in a compatible vector model. In addition, a study looking at transcriptional changes in both *B. malayi* and *A. aegypti* during the course of infection saw that between 2-4 days post infection, the *A. aegypti* serine protease gene, AAEL002590, was downregulated in infected mosquitoes (84). This downregulation occurring 2-4 days post infection correlates with our study as 2 days post infection broadly aligns with the mf to L1 molt within the thoracic muscles but some mf will still be present (43). It is also important to note that while our data shows downregulation occurring as early as 24 hours post treatment, this may be due to the fact that we are injecting isolated EVs and not infecting with live parasites. Many of these transcriptomic studies also identified downregulation of CLIP serine proteases or prophenoloxidase enzymes (43,83,85), key components involved in the prophenoloxidase cascade and melanization immune response, providing additional corroboration for our observation that *B. malayi* mf EVs are interfering with this immune response. While our study provides a mechanism for parasite-derived transcriptional changes of the host, all these transcriptional studies provide a substrate for further studies aimed at better understanding mosquito immune responses. Paying particular attention to those mosquito genes that filarial nematode parasites have been selected to suppress over millions of years of the host-parasite interaction may reveal the most critical pathways and proteins to exploit for insecticides or novel transmission control strategies.

There is a growing body of evidence that host immunomodulation by parasite-derived EVs is a common motif in parasitic nematode infections. This picture began to emerge with seminal work from Buck et al. (2014) who showed that EVs released by the murine gastrointestinal nematode, *Heligmosomoides polygyrus* suppressed expression of an IL-33 receptor subunit (also known as ST2) in intestinal epithelial cells (15, 16). IL-33 is an alarmin cytokine that plays an important role in initiation of type 2 immune responses, is critical for driving induction of Th2-associated cytokines, and is involved in the expulsion of intestinal parasitic nematodes (86). This work was extended to show that the same EVs elicited similar modulatory phenotypes in macrophages (16). Further studies have shown that *Trichinella spiralis* EVs are capable of producing some of the modified type 2 immune response phenotypes seen in chronic infections. *T. spiralis* EVs have been shown to downregulate the pro-inflammatory cytokines IL-1β, TNFα, and IFNγ while also increasing production of the anti-inflammatory cytokines IL-10 and TGF-β in an induced colitis mouse model (4). Similarly, *Nippostrongylus brasiliensis* EVs were able to reduce IL-1β and increase IL-10 expression in a similar induced colitis mouse model (55). Importantly, our group and others have shown that the modulation of host biology via EVs is not limited to gastrointestinal parasitic nematodes but also occurs at the filarial nematode-host interface. We have previously described how EVs released by infective L3 stage *B. malayi* drive a phenotype in murine macrophages that is more consistent with classical activation than alternative activation (44). More compelling, evidence generated by the Nutman laboratory shows that EVs secreted by *B. malayi* mfs inhibit phosphorylation of the mTOR complex in human monocytes and point to these EVs as the critical parasite-derived factor eliciting Dendritic cell dysfunction during filarial disease (53). The data from these various studies collectively show parasite-derived EVs driving diverse but consistent effects in mammalian host, an observation that we now extend to the vector host.

Our understanding of the filarial nematode-vector interface is incomplete, but the data described in this paper helps to begin addressing this knowledge gap and may even seed the identification of novel targets that could contribute to better controlling filarial nematode diseases. Identifying targets at the vector stage of parasite development may stop transmission of the causative agents of filarial diseases and may provide insight into control strategies for other non-filarial, vector-borne diseases. Any mechanism that disrupts the vector-parasite interaction and skews the balance in favor of the vector is likely to prevent infection, parasite development and transmission.

## 4. Materials and Methods

### 4.1 Cell culture

The immunocompetent *Aedes aegypti*-derived Aag2 cell line was cultured in Schneider’s *Drosophila* medium supplemented with 10% heat-inactivated, fetal bovine serum and 1% Penicillin/Streptomycin (all Thermo Fisher Scientific, Waltham, MA, USA) at 28°C.

### 4.2 Parasite Culture and Maintenance

*Brugia malayi* parasites were obtained from the NIH/NIAID Filariasis Research Reagent Resource Center (FR3) at the University of Georgia, USA. Persistent *B. malayi* infections at FR3 are maintained in domestic short-haired cats. Microfilariae stage *B. malayi* were obtained from a lavage of the peritoneal cavity of a euthanized gerbil. Microfilaria were washed according to FR3 protocols upon arrival at Iowa State University. Briefly, microfilariae were centrifuged at 2000 rpm for 10 minutes at room temperature to pellet parasites. Transport media [RPMI with Penicillin (2000 U/ml) and Streptomycin (2000 µg/ml)] was aspirated and the parasite pellet resuspended in dPBS (Thermo Fisher Scientific). The parasite suspension was overlaid onto 10 ml of Histopaque-1077 (Sigma Aldrich, St. Louis, MO, USA) and centrifuged at 2000 rpm for an additional 15 minutes. The supernatant was aspirated and parasite pellet washed with dPBS twice for 5 minutes each wash. After washing, the supernatant was aspirated and 3 ml of cell culture grade water (Cytiva, Marlborough, MA, USA) was added to the remaining pellet to lyse red blood cells (RBCs). Immediately following RBCs lysis, 10 ml dPBS was added and parasites centrifuged for an additional 5 minutes then washed one final time in dPBS. Microfilariae were then resuspended in worm culture media (RPMI with 1% 1 M HEPES, 1% 200mM L-glutamine, Penicillin (2000 U/ml), Streptomycin (2000 µg/ml), and 1% w/v glucose [all Thermo Fisher Scientific]) and cultured at 37°C with 5% CO_2_ for 5-7 days. Parasite motility was used as an indicator of parasite viability. Parasite viability was checked daily and spent media was collected every 24 hours and retained for EV isolation as long parasites appeared viable.

### 4.3 Mosquito Rearing

*A. aegypti* (Liverpool strain) mosquitoes were reared at 27°C and 80% relative humidity with a 14:10 h light/dark period. Larvae were fed a 50:50 diet of Tetramin ground fish flakes (Tetra, Melle, Germany) and milk bone dog biscuits. Adults were maintained on a 10% sucrose solution. All experimental techniques were performed on cohorts of 4–6 days old adult female mosquitoes.

### 4.4 EV Isolation, Quantification & Imaging

EVs were isolated from spent culture media via differential ultracentrifugation as previously described (44,46,47). Briefly, media was filtered through 0.2 μm PVDF filtered syringes (GE Healthcare, Chicago, IL, USA) and centrifuged at 120,000 x *g* for 90 minutes at 4°C. The supernatant was decanted leaving approximately 1.5 ml media to ensure that the EV pellet was not disrupted. The retained media and pellet were filtered through a PVDF 0.2 μm syringe filter and centrifuged at 186,000 x *g* for a further 2 h at 4°C. The size profile and concentration of EVs in the isolated sample were quantified using nanoparticle tracking analysis (NTA; NanoSight LM10, Malvern Instruments, Malvern, UK). EV integrity and morphology were confirmed using transmission electron microscopy (TEM). Briefly, a 2 µl aliquot of EV preparation was placed onto a carbon film grid (Electron Microscopy Sciences, Hatfield, PA, USA) for 1 minute. The drop was wicked to a thin film and 2 µl of uranyl acetate (2% w/v final concentration) was immediately applied for 30 seconds, wicked, and allowed to dry. Images were taken using a 200kV JEOL 2100 scanning and transmission electron microscope (Japan Electron Optics Laboratories, LLC, Peabody, MA) with a Gatan OneView camera (Gatan, Inc. Pleasanton, CA).

### 4.5 EV internalization by Aag2 cells

Methods were based on protocols previously described (46), but modified for optimal imaging of the Aag2 cell line. 3 ×10^5^ Aag2 cells were seeded on an 18 mm, #1 thickness, poly-D-lysine coverslip (Neuvitro, Vancouver, WA) in a 12-well plate (Thermo Fisher Scientific) and cultured at 28°C overnight. Between 5×10^8^-1×10^9^ isolated EVs were stained with PKH67 (Sigma Aldrich, St. Louis, MO) according to manufacturer’s instructions. Confluent Aag2 cells were treated with 3.5×10^7^ stained EVs and incubated for 24 hrs at 28°C. EV uptake was visualized with immunocytochemistry. Media was removed and cells were washed with 1X dPBS and fixed in 4% paraformaldehyde (Electron Microscopy Sciences) for 15 minutes at room temperature. Following three 1X dPBS washes at room temperature, cells were incubated with 1:300 Alexa Fluor 647 phalloidin (Thermo Fisher Scientific) for 45 minutes at room temperature followed by three washes of 1x dPBS for 5 minutes each. Cells were incubated with 300 nM DAPI (Thermo Fisher Scientific) for 5 minutes at room temperature followed by two washes in 1X dPBS. Coverslips were mounted using Flouromount aqueous mounting media (Sigma Aldrich) and visualized by a Leica SP5 X MP confocal/multiphoton microscope system (Leica Microsystems Inc., Buffalo Grove, IL, USA).

Concurrently, EV internalization was quantified using flow cytometry. 3 ×10^5^ cells were seeded per well of a 12-well plate and incubated at 28°C overnight. Cells were incubated with 3.5×10^7^ PKH67 stained EVs for 24 hrs at 28°C. Cells were washed in 1x dPBS and collected into polystyrene FACS tubes (Thermo Fisher Scientific). Cells were fixed in 4% paraformaldehyde for 20 minutes and washed with FACS buffer (dPBS supplemented with 1% BSA and 0.1% NaN_3_). Cells were resuspended in 400 µl FACS buffer and analyzed with a BD Accuri C6 Flow Cytometer (BD Biosciences, San Jose, CA). For endocytosis inhibition assays, Aag2 cells were treated with a final concentration of either 30 µM chlorpromazine or 15 µM nystatin (Thermo Fisher Scientific). Following a two-hour incubation, media was changed and cells treated with 3.5×10^7^ *B. malayi* mf EVs, incubated for 24 hours and then collected for confocal microscopy and flow cytometry as described above.

### 4.6 mRNA-Seq Analysis

1 x 10^5^ Aag2 cells were seeded in each well of a 96-well plate (Corning Inc, Corning, NY, USA) and incubated overnight at 28°C. The following day, culture media was changed and cells were treated with either lipopolysaccharide (LPS) (500 ng/ml) to stimulate an immune response *in vitro* or dPBS as a negative control. Cells were incubated for an additional 12 hours at 28°C after which, culture media was changed and cells treated with 1.1 x 10^9^ parasite EVs per well. Cells were then incubated for a further 16 hours at 28°C before collection and storage in Trizol (Thermo Fisher Scientific) ahead of RNA extraction. Briefly, cells in Trizol were mixed with chloroform (0.2 ml chloroform per ml Trizol) and shaken vigorously for 20 seconds. Samples were allowed to sit at room temperature for 3 minutes and then centrifuged at 10,000 x *g* for 18 minutes at 4°C. The aqueous phase was collected, and an equal volume of 100% ethanol was added. RNA was then purified and collected using a RNeasy Mini Kit (Qiagen, Hilden, Germany) according to manufacturer’s instructions.

mRNA-seq was performed by LC Sciences (Houston, TX). Total RNA quantity and purity were analyzed using an RNA 6000 Nano LabChip Kit and a Bioanalyzer 2100 (Agilent, Santa Clara, CA). High quality RNA samples with RIN number > 7 were used to construct the sequencing library. mRNA was purified from total RNA (5µg) using Dynabeads Oligo (dT)(Thermo Fisher Scientific) with two rounds of purification. Following purification, mRNA was fragmented into short fragments using a NEB Next Magnesium RNA Fragmentation Module (New England Biolabs, Ipswich, MA, USA) at 94°C for 5-7 minutes. Cleaved RNA fragments were reverse transcribed to cDNA by Superscript II Reverse Transcriptase (Thermo Fisher Scientific) and the resulting cDNA used to generate U-labeled second-stranded DNA using *E. coli* DNA polymerase I, RNase H (both New England Biolabs) and dUTP Solution (Thermo Fisher Scientific). An A-base was added to the blunt ends of each strand, preparing them for ligation to the indexed adapters. Each adapter contained a T-base overhang for ligating the adapter to the A-tailed fragmented DNA. Dual-index adapters were ligated to the fragments, and size selection was performed with AMPureXP beads (Beckman Coulter, Brea, CA, USA). U-labeled second-stranded DNAs were treated with heat-labile UDG enzyme (New England Biolabs), and ligated products were amplified with PCR by the following conditions: initial denaturation at 95°C for 3 minutes; 8 cycles of denaturation at 98°C for 15 seconds, annealing at 60°C for 15 seconds, and extension at 72°C for 30 seconds; and final extension at 72°C for 5 minutes. The average insert size for the paired-end libraries was 300 bp (±50 bp). Paired-end sequencing was performed on an Illumina Hiseq 4000 (Illumina, San Diego, CA, USA). Reads were adapter and quality trimmed using Trimmomatic (87). HISAT2 (88) and StringTie (89) were used to align surviving reads to the *B. malayi* reference genome (WormBase ParaSite version 12.4) (90, 91) and to the *A. aegypti* reference genome (VectorBase release 47) (92) to produce raw counts for annotated genes. The RNA-seq pipeline was implemented using Nextflow (93). DESeq2 (94) and custom R scripts were used to identify differentially expressed genes (DEGs) across conditions. The R package topGO (95) was used to assess functional enrichment of differentially expressed genes. Gene ontology (GO) terms from the *A. aegypti* LVP transcriptome were retrieved from VectorBase (92).

### 4.7 miRNA-Seq Analysis

microRNA (miRNA) sequencing was performed by LC Sciences. The total RNA quality and quantity were analyzed by Bioanalyzer 2100 (Agilent Technologies, Santa Clara, CA) with RIN number >7.0. Small RNA libraries were prepared using 1 µg of total RNA and the TruSeq Small RNA Sample Prep Kits (Illumina) according to manufacturer’s instructions. Single-end sequencing was performed on an Illumina Hiseq 2500 (Illumina) according to manufacturer’s instructions. Raw reads were subjected to an in-house program, ACGT101-miR (LC Sciences), to remove adapter dimers and junk, low complexity and common non-target RNA families (rRNA, tRNA, snRNA, snoRNA) and repeats. Remaining unique sequences with length 18∼26 nucleotides were mapped to specific species precursors in miRBase 22.0 (96–101) and by BLAST search (102) to identify known miRNAs and novel 3p- and 5p-derived miRNAs with their genomic location. Length variation at both 3’ and 5’ ends and one mismatch inside of the sequence were allowed in the alignment. The unique sequences mapping to specific species mature miRNAs in hairpin arms were identified as known miRNAs. The unique sequences mapping to the other arm of known specific species precursor hairpin opposite to the annotated mature miRNA-containing arm were considered to be novel 5p- or 3p derived miRNA candidates. Hairpin RNA structures of unmapped sequences were predicted from the flanking 80 nucleotide sequences using RNAfold (103). The criteria for secondary structure prediction included number of nucleotides in one bulge in stem (≤12), number of base pairs in the stem region of the predicted hairpin (≥16), cutoff of free energy (kCal/mol ≤-15), length of hairpin (up and down stems + terminal loop ≥50), length of hairpin loop (≤20), number of nucleotides in one bulge in mature region (≤8), number of biased errors in one bulge in mature region (≤4), number of biased bulges in mature region (≤2), number of errors in mature region (≤7), number of base pairs in the mature region of the predicted hairpin (≥12) and percent of mature sequences in stem (≥80). To predict the genes targeted by most abundant miRNAs, two computational target prediction algorithms TargetScan (104–106) and Miranda 3.3a (107) were used to identify putative miRNA binding sites. Finally, the data predicted by both algorithms were combined and the overlaps calculated. The R package, enrichplot, was used to visualize GO term enrichment from the predicted targets of differentially expressed miRNAs.

### 4.8 RT-qPCR Validation of Gene Expression Levels

1×10^5^ Aag2 cells were seeded in each well of a 96-well plate (Corning Inc, Corning, NY, USA) and incubated overnight at 28°C. The following day, culture media was changed and cells were treated with either LPS (500 ng/ml) to stimulate an immune response *in vitro* or dPBS as a negative control. Cells were incubated for an additional 12 hours at 28°C after which, culture media was changed and cells treated with 10-fold serial dilutions ranging from 1 x 10^9^ to 1 x 10^2^ parasite EVs. This range was used as it allowed us to see the effects of treating cells with more EVs than would be present in a natural infection, EV levels present during a natural infection (1 x 10^6^ – 1 x 10^5^) and the effects of having less EVs than would occur in a natural infection. Cells were then incubated for a further 16 hours at 28°C before collection and storage in Trizol (Thermo Fisher Scientific) ahead of RNA extraction described above. cDNA was synthesized from sample RNA using Superscript III First-Strand cDNA Synthesis kit (Thermo Fisher Scientific) according to manufacturer’s instructions. 20 ng of cDNA was used per qPCR reaction using Powerup SYBR green master mix (Thermo Fisher Scientific) and gene specific primers according to manufacturer’s instructions on a Quantstudio 3 Real-Time PCR system (Thermo Fisher Scientific). CT values were averaged across technical replicates and normalized against RPS17. Primer sequences for AAEL002590 (Serine Protease), AAEL024490 (predicted cys-loop ligand-gated ion channel [cysLGIC] subunit), and the housekeeping gene (RPS17) can be found in Supplemental Table 1.

### 4.9 *In vitro* RNA Interference

Duplexed siRNA was designed and produced targeting the serine protease gene by Integrated DNA Technologies (Coralville, IA, USA). Sequences for the duplexed siRNA can be found in Supplemental Information 3. 4 x 10^4^ Aag2 cells were seeded per well of a 96-well plate and incubated overnight. 5 pmol of siRNA or scrambled negative control was mixed with lipofectamine RNAiMAX Reagent (Thermo fisher Scientific) to create a 1 pmol siRNA solution. 10 µl of the 1 pmol siRNA solution was added per well and incubated for 24 hours. To determine RNAi efficiency, total RNA was isolated from cells for subsequent RT-qPCR as described above.

### 4.10 *Aedes aegypti* Injections

Four to five-day old *A. aegypti* (Liverpool strain) female mosquitoes were intrathoracically injected with 69 nl of LPS (1 mg/ml) [Sigma Aldrich] or dPBS (Thermo Fisher Scientific) using a Nanoject III injector (Drummond Scientific Company, Broomall, PA, USA) and incubated for six hours at 27°C prior to EV injection. Mosquitoes were then challenged with serial dilutions of 1×10^7^, 1×10^6^, 1×10^5^ EVs, or dPBS as a control. Total RNA was isolated from 8 mosquitoes per treatment group 24 hours post-challenge. Mosquitoes were homogenized using a mortar and pestle in 1 ml of Trizol. The resulting suspension was centrifuged at 12,000 x *g* for 10 minutes at 4°C to remove debris, the supernatant collected. RNA extraction, cDNA synthesis and qPCR were performed as previously described.

### 4.11 Phenoloxidase Activity Assay

Pooled hemolymph was collected from 10 adult female mosquitoes by perfusion and prepared for PO assay as previously described (108). Briefly, 10 µl of hemolymph was mixed with 90 µl of 3, 4-Dihydroxy-L-phenylalanine (L-DOPA, 4 mg/ml)(Sigma Aldrich) dissolved in nuclease free water (Cytiva). After an initial 10 minutes incubation at room temperature, PO activity was measured at 490 nm every 5 minutes for 30 minutes, then the final activity was measured at 60 minutes using a Synergy HTX Multi-Mode Microplate Reader (Agilent). To determine if AAEL002590 was directly involved in the PO pathway AAEL002590 was knockdown via siRNA in Aag2 cells as previously described. After the 24 hr incubation, Aag2 cells were challenged with LPS (500ng/ml) for either 6 or 24 hours. 10 µl of either control or siRNA treated Aag2 cell culture media was mixed with 90 µl L-DOPA as previously described. PO activity was allowed to proceed at room temperature overnight after which PO activity was measured at 490 nm.

### 4.12 Statistical Analysis

*In vitro* RT-qPCR validation was analyzed using a repeated measures one-way ANOVA while *in vivo* RT-qPCR validation was analyzed using mixed effects one-way ANOVA. Multiple comparisons were conducted using the Dunnett statistical hypothesis testing method. Enrichment of functions within the molecular function, biological process, and cellular component GO term sub-ontologies were analyzed using a Fisher’s exact test. *In vivo* PO assays were analyzed using a repeated measures two-way ANOVA with a Šidák multiple comparisons test while in vitro PO assays were analyzed with multiple T tests with a Holm-Šidák multiple comparison test. For all significance testing p-values < 0.05 was considered significant. All ANOVAs were completed using GraphPad prism 9.3.1 (GraphPad Software, San Diego, CA, USA).

