## Supplemental Materials for "Extracellular vesicles secreted by *Brugia malayi* microfilariae modulate the melanization pathway in the mosquito host"

| Experiment | Gene | Forward Primer | Reverse Primer |
|---|---|---|---|
| RT-qpcr Gene Expression Validation | | | |
| | AAEL002590 | AGAGCCTAGTTGCGTTGTTAG | CTGTACTGACTTCTGTGGGAAC |
| | AAEL024490 | GCGACACCTCATCCTTTCTT | CTCGTCGTTCTTCCAGACATAC |
| | Housekeeping Gene (RPS17) | CACTCCCAGGTCCGTGGTAT | GCACACTTCCGGCACGTAGT |
| RNAi | | | |
| | Duplexed siRNA for AAEL002590 | AAAGAUAUCAUUGCUAGUGACCAAA | UCUUUCUAUAGUAACGAUCACUGGUUU |

**Supplemental Table 1: Primer Sequences**

Primer sequences for RT-qpcr validation of AAEL002590 (serine protease) and AAEL024490 (CLIC subunit). RT-qpcr data was normalized against the housekeeping gene RPS17. Duplexed siRNA for knockdown of AAEL002590 was produced by Integrated DNA Technologies with the sequences above.

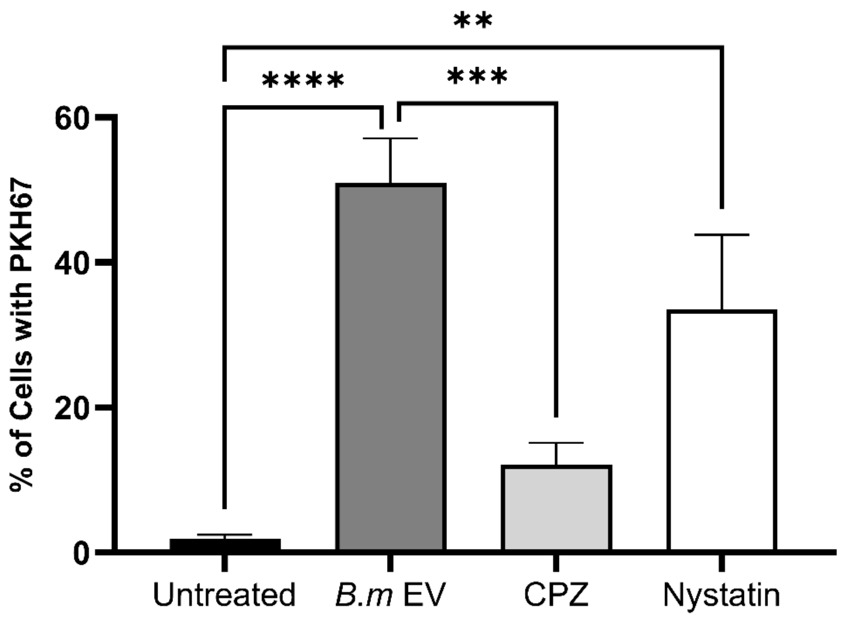

**Supplemental Figure 1: Quantification of EV Internalization in Aag2 cells**

EV internalization by Aag2 cells treated with or without *Brugia malayi* microfilariae (mf) extracellular vesicles (EVs) was quantified using a BD Accuri C6 Flow Cytometer. 51% of Aag2 cells internalized *B. malayi* mf EVs as compared to untreated control. The endocytosis inhibitors chlorpromazine (CPZ) and nystatin were used to determine the mechanism by which these EVs are being internalized. CPZ significantly reduced EV internalization by 24% as compared to EV treated cells while nystatin did not significantly reduce EV internalization as compared to EV only treated cells. N = 3 (minimum). Mean ± SEM. **$P < 0.01$, ***$P < 0.001$, ****$P < 0.0001$.

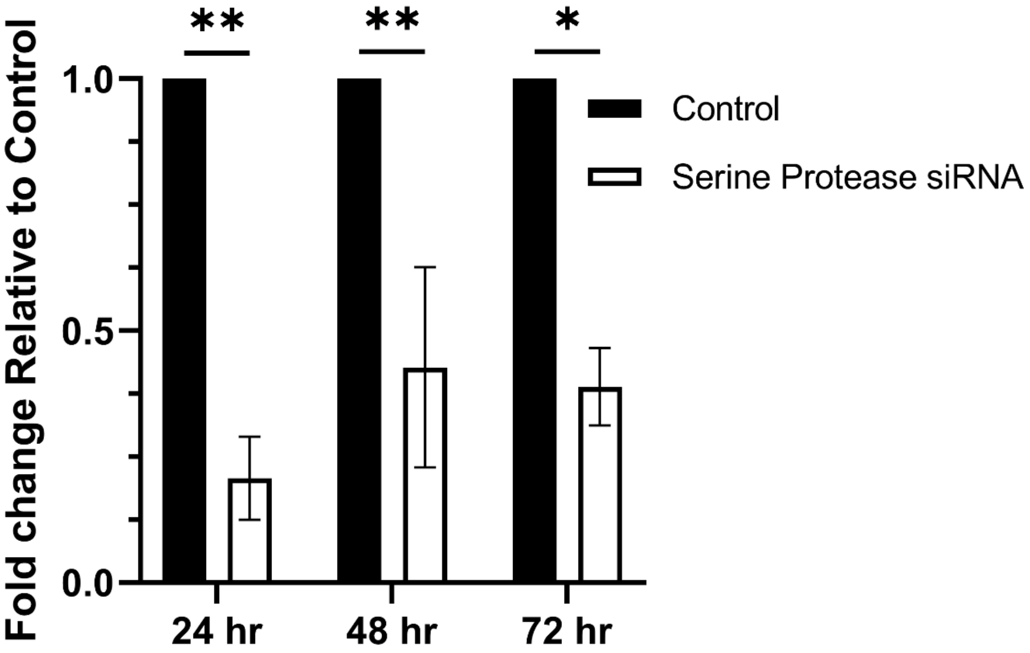

**Supplemental Figure 3: Efficiency time course of RNAi knockdown of AAEL002590**

Aag2 cells were treated with 1 pmol (final concentration) of duplexed siRNA. Gene expression of AAEL002590 was quantified using RT-qpcr. Expression of AAEL002590 was significantly reduced 79%, 57%, and 61% at 24 hrs, 48 hrs and 72 hrs post-treatment, respectively, as compared to control cells. Knockdown was the most efficient 24 hrs post-treatment therefore this time point was used in all experiments going forward. N = 3 (minimum). Mean ± SEM. *P < 0.05, **P < 0.01.
